# ProtPipe2: Multi-Platform Downstream Proteomics Analysis Tool

**DOI:** 10.64898/2026.09.15.751860

**Authors:** Jacob Epstein, Cory Weller, Isabelle Kowal, Ying Hao, Cole Tindall, Benjamin Jin, Mark R. Cookson, Mike A. Nalls, Ziyi Li, Yue A. Qi

**Affiliations:** Center for Alzheimer’s and Related Dementias (CARD), National Institute on Aging and National Institute of Neurological Disorders and Stroke, National Institutes of Health, 9000 Rockville Pike, Bethesda, Maryland 20892, USA; DataTecnica LLC, 1275 25th Street NW Suite 805, Washington, DC 20037, USA; Cell Biology and Gene Expression Section, Laboratory of Neurogenetics, National Institute on Aging, National Institutes of Health, 9000 Rockville Pike, Bethesda, Maryland 20892, USA

**Keywords:** proteomics, R, Shiny, mass spectrometry, Olink, SomaScan

## Abstract

Proteomics studies increasingly use different technologies, including mass spectrometry, the antibody-based Olink platform, and the aptamer-based SomaScan platform. However, downstream analysis often depends on platform-specific scripts or point-and-click tools, limits flexibility and makes analyses difficult to reuse across platforms. We developed ProtPipe2, a downstream proteomics analysis framework for mass spectrometry, Olink, and SomaScan data that is available as both an R package and an interactive web application. ProtPipe2 supports quality control, preprocessing, imputation, batch correction, statistical analysis, visualization, pathway analysis, and result export. ProtPipe2 expands the original Protpipe workflow to additional proteomics platforms and provides reusable R functions for imputation, batch correction, and other downstream analyses. Its redesigned web application provides user-controlled methods and parameters through the same R-based workflow. Case studies across mass spectrometry, Olink, and SomaScan datasets demonstrated a consistent downstream analysis workflow across platforms. By supporting three proteomics platforms through both graphical and scripted workflows, ProtPipe2 makes downstream analysis easier to perform, repeat, and review. ProtPipe2 is freely available at https://nih-card.github.io/ProtPipe2/.

## INTRODUCTION

Proteomics is no longer defined by a single measurement technology. Mass spectrometry supports broad protein discovery, whereas Olink and SomaScan enable large-scale protein measurement using antibodies and aptamers, respectively. Although these technologies differ in their measurement methods and output formats, they share many downstream analytical needs, including sample-level quality control, missing-value handling, normalization, batch assessment, statistical analysis, and pathway interpretation.

Mass spectrometry, Olink, and SomaScan data differ in scale, measurement units, quality flags, detection-limit information, and metadata structure.^1,2^ Researchers must therefore often combine platform-specific import scripts with separate tools for preprocessing, statistical analysis, and visualization. This fragmented approach makes analytical choices harder to track and workflows harder to reproduce. Graphical tools improve accessibility, but analyses performed through an interface may remain difficult to repeat in code or adapt to another study.

Existing software addresses parts of this workflow. The original ProtPipe combines data-independent-acquisition processing and downstream reports in a command-line and container-oriented workflow with a separate web interface.^3^ Perseus provides a mature desktop environment for label-based and label-free MS data;^4^ amica accepts processed proteomics tables and adds quality control, differential expression, and over-representation analysis;^5^ ProTIGY offers interactive exploration of any quantitative feature-by-sample matrix;^6^ and MSstatsShiny provides a graphical interface to the MSstats statistical framework.^7^ These tools are valuable, but the evaluated versions primarily focus on mass spectrometry data or require a previously prepared generic abundance matrix. Researchers working with long-form Olink NPX exports or SomaScan ADAT files may therefore need additional platform-specific processing before beginning a general downstream analysis.

ProtPipe2 was developed to provide a common downstream workflow for mass spectrometry, Olink, and SomaScan proteomics data. It is available as an R package and an interactive web application, allowing users to perform the same core analyses through either scripted or graphical workflows. Here we describe the architecture and analysis assumptions, compare the design with the original ProtPipe and related software, demonstrate three platform-specific inputs, and measure package and application scaling. The case studies demonstrate software behavior and do not claim independent biological replication of the source publications.

## METHODS

### Software Implementation and Data Representation

ProtPipe2 is implemented as an open-source R package with a companion application developed in Shiny. The source code is available at https://github.com/NIH-CARD/ProtPipe2, and package documentation is available at https://nih-card.github.io/ProtPipe2/.

All input types are converted to a `SummarizedExperiment` object^8^, which stores protein or assay measurements in an assay named *intensities*, feature annotations in `rowData`, sample annotations in `colData`, and analysis provenance in `metadata`. Because every package function receives and returns this object, the same data move freely between scripted and interactive analyses without conversion to a separate internal format. The Shiny application calls the package functions directly rather than maintaining an independent analysis implementation.

Each preprocessing function appends its operation name, parameter values, and processing details to an ordered log stored in the object metadata, and this complete history can be exported as a Markdown report. Sample identifiers are converted to unique, syntactically valid R names; consequently, identifiers beginning with a number receive an X prefix in the analysis object and in exported results.

### Generic Matrix Import and Metadata Validation

Generic input consists of a protein-by-sample abundance table and an optional sample-metadata table. The Shiny application accepts CSV, TSV, XLS, and XLSX files, whereas the R interface accepts a data frame. Protein annotations occupy the non-intensity columns, and intensity columns can be selected explicitly. When no intensity columns are specified, numeric columns are interpreted as measurements; explicit selection is therefore recommended whenever the feature annotations contain numeric variables.

The sample-metadata table is required to contain a SampleID column, and duplicate metadata identifiers halt import. Metadata identifiers absent from the abundance matrix are removed with a warning, whereas assay samples lacking matching metadata are retained with missing metadata values. Metadata rows are then reordered to match the assay columns. When no metadata table is supplied, ProtPipe2 generates basic sample annotations from the abundance- column names.

### Platform-Specific Import

For Olink data, long-form NPX files are read with `OlinkAnalyze` and processed using the ProtPipe2 Olink import functions. Samples are excluded if any corresponding `QC_Warning` value is missing or is not recorded as *PASS*. Assay-level warning flags and NPX measurements below the supplied limit of detection are converted to missing values. The filtered long-form data are then pivoted to a feature-by-sample matrix; when more than one value is present for the same assay and sample, the values are represented by their mean. UniProt identifiers, assay names, Olink identifiers, and detection-limit information are retained as feature annotations. Because NPX is already reported on a log2 scale, the imported object is marked as log2 transformed, preventing an unintended second transformation.

SomaScan ADAT files are read using *SomaDataIO*^9^. Rows identified as study samples are separated from buffer and calibrator rows. All study samples with duplicated `SampleId` values are excluded. Assay measurements are extracted from the SomaScan sequence columns, and analyte annotations—including the assay name, UniProt identifier, gene symbol, target name, organism, and assay type—are retained in `rowData`. Feature-specific buffer and calibrator means are also retained as feature annotations. When buffer filtering is selected, study-sample measurements below the corresponding feature-specific mean buffer value are converted to missing values. ProtPipe2 uses the vendor-processed ADAT measurements and does not repeat SomaScan hybridization normalization, median-intensity normalization, plate scaling, calibration, or vendor quality-control procedures.

### Quality Control

Quality-control functions calculate the number of quantified features per sample, sample-intensity distributions, within-group protein coefficients of variation, pairwise sample correlations, and feature- and sample-level missing-value rates. Correlations use Spearman rank correlation with pairwise-complete observations and may be calculated from all proteins or from a specified number of the highest-variance proteins. The web application initially selects the 1,000 highest-variance proteins for the correlation display, although users may select another number or use all features.

The definition of a detected measurement depends on the platform. For processed MS matrices, non-missing measurements are considered detected. For Olink, samples with sample-level QC warnings are first excluded, and a measurement is considered detected when its NPX value is over the assay- and plate-specific limit of detection. For SomaScan, a measurement is considered detected when its RFU value is above the corresponding aptamer-specific buffer value. ProtPipe2 uses these definitions to calculate the percentage of detected measurements for each protein and sample. Quality-control calculations and plots do not remove proteins or samples and do not alter the reported abundance values.

### Pre-processing

ProtPipe2 organizes preprocessing into six steps: minimum-intensity filtering, protein and sample filtering, transformation, normalization, imputation, and batch-effect assessment and correction. Each operation is optional, and the selected parameters and numbers of removed proteins or samples are recorded in the processing history.

### Minimum-intensity filtering

Minimum-intensity filtering is available for processed MS data. When enabled, measurements below a user-defined intensity are classified as undetected and replaced with missing values before protein- and sample-level filtering. This option is not applied to Olink or SomaScan data, for which detection is determined using the platform-specific references described in the Quality Control section.

### Protein and sample filtering

Protein and sample filtering follows the same general workflow for MS, Olink, and SomaScan data. For MS data, detection is based on nonmissing measurements after any minimum-intensity filtering. For Olink, detection requires an NPX value at or above the assay- and plate-specific LOD without an assay-level QC flag. For SomaScan, detection requires an RFU value at or above the aptamer-specific buffer value. Proteins are first retained according to a user-defined detection rate across samples. Sample detection rates are then recalculated using the retained proteins, followed by sample filtering. The web application initially displays thresholds of 80% for proteins and 95% for samples, but filtering remains disabled until selected by the user.

### Transformation and normalization

The optional transformation step calculates log2(x + 1). Additional normalization is also optional. Median normalization rescales each sample to the global median, whereas mean normalization rescales each sample to the global mean. The web application initially displays median normalization, but normalization remains disabled until selected. Additional normalization of Olink or vendor-processed SomaScan data should be performed only when supported by the study design.

### Imputation

ProtPipe2 provides several optional methods for imputing genuinely missing abundance values, including fixed-value replacement, scaled protein-minimum replacement, protein-wise mean or median replacement, a protein-wise left-shifted normal distribution, K-nearest-neighbor imputation, and random-forest imputation. The choice of method should reflect the expected missingness mechanism. Left-shifted imputation is intended for MS measurements expected to be missing because of low abundance, whereas K-nearest-neighbor and random-forest methods use relationships among proteins or samples to estimate missing values. Mean and median imputation provide simple alternatives but can reduce the observed variability of a protein.

No imputation is applied by default. For MS data, the web application identifies left-shifted imputation as a suggested starting method when the missing values are consistent with left-censored, low-abundance measurements. The starting mean shift is 1.8 standard deviations, and the starting distribution width is 0.3 standard deviations. K-nearest-neighbor imputation initially uses 10 neighbors. Random-forest imputation is presented as an advanced, computationally intensive option. Random seeds are recorded for stochastic methods.

For Olink and SomaScan data, no imputation is recommended by default. Measurements below the Olink LOD or SomaScan buffer reference remain available as reported values and are not converted to missing values. Users may enable an imputation method only when genuine missing values remain and the selected method is supported by the study design. All imputation choices and parameter values are recorded in the processing history.

### Batch-effect assessment and correction

Batch correction is never applied automatically. When explicitly selected, correction is performed using limma::removeBatchEffect.^10^ Users specify the batch variable and any biological variables whose associated variation should be retained. Correction is stopped if any sample lacks a batch assignment. The batch variable, protected biological variables, and correction settings are recorded in the processing history.

The values displayed in the web application are starting settings rather than universal analytical recommendations. Appropriate filtering thresholds, transformation, normalization, imputation, and batch-correction settings depend on the measurement platform, experimental design, data scale, and expected missingness mechanism. The platform availability, starting values, and user-adjustable settings for all preprocessing operations are summarized in Supplementary Table 1.

### Clustering and Dimensionality Reduction

Hierarchical clustering is performed across samples using Euclidean distance and complete linkage by default. Principal component analysis is performed using centered and scaled measurements after near-zero-variance features are removed. UMAP is calculated with the R `umap` package, using 15 neighbors as the package-function default; the neighbor number is reduced when required by the sample count. The Shiny application sets a random seed before UMAP calculation. Hierarchical clustering, principal component analysis, and UMAP require complete measurements, so missing values suggested to be removed or imputed before these analyses.

### Statistical Analysis

Two-group differential-abundance analysis is performed using `limma`.^10^ If an input object has not been identified as log2 scaled, ProtPipe2 applies log2(x + 1) before model fitting. The software constructs a cell-means design matrix and can incorporate user-specified covariates. The requested treatment-versus-reference contrast is evaluated using `lmFit`, `contrasts.fit`, and empirical-Bayes moderation with `eBayes(trend = TRUE)`. Complete measurements and at least two samples in each comparison group are required.

For categorical variables, ProtPipe2 performs moderated group comparisons to identify proteins that differ between two or more groups. For continuous variables, such as age, clinical measurements, or quantitative scores, ProtPipe2 evaluates the association between each protein and the selected variable using Spearman rank correlation. P values are adjusted across proteins using the Benjamini–Hochberg procedure. Nominal P-value and fold-change thresholds used to highlight figures are reported as visualization criteria and are distinguished from false-discovery-rate-controlled findings.

### Pathway Analysis

Gene symbols are mapped to Entrez Gene identifiers using `clusterProfiler::bitr`.^11^ When an annotation contains multiple semicolon-delimited symbols, the first symbol is used for mapping. Gene Ontology over-representation analysis uses all successfully mapped proteins measured by the assay as the background universe rather than using the complete genome. Enrichment P values are adjusted using the Benjamini–Hochberg procedure.

Gene-set enrichment analysis ranks mapped proteins by log2 fold change for categorical comparisons or by Spearman correlation coefficient for continuous associations. Gene Ontology, KEGG, and user-supplied gene sets are supported, with custom gene sets accepted in GMT, CSV, or TSV format. Because the multi-group omnibus test does not yield a single directional effect across all groups, the Shiny workflow restricts multi-group results to over- representation analysis unless a directional contrast is defined separately.

### Case Study Data and Analysis

#### Mass spectrometry

The bundled `iPSC.csv` matrix contained 9,119 protein rows and 42 samples across seven time points from day 0 to day 28. The generic-matrix workflow was used to calculate within-time-point coefficients of variation, Spearman sample correlations, hierarchical clustering, principal components, and targeted protein heatmaps. Missing measurements used in analyses requiring complete data were replaced with the specified fixed value.

#### Olink

Long-form NPX data and the associated sample manifest were obtained from PRIDE Affinity Proteomics accession PAD000040. The input contained 16,284 measurements from 142 samples and 92 assays distributed across two inflammation-panel plates. Sixteen samples carrying QC warnings were excluded, leaving 126 samples. Detection status was calculated for each measurement using its plate-specific LOD and assay-level flag before repeated sample–assay measurements were combined. Assays detected in at least 80% of the 126 samples were retained first, followed by samples detected for at least 95% of the retained assays. This produced a matrix of 74 assays and 124 samples: 59 without diabetes, 22 with prediabetes, and 43 with type 2 diabetes. Reported NPX values, including below-LOD values belonging to retained assays and samples, were preserved, and no imputation was applied. Moderated two-group models compared prediabetes and type 2 diabetes separately with the no-diabetes reference. Nominal P ≤ 0.05 and |log2 fold change| ≥ 0.25 were used only to highlight the volcano plots. A moderated omnibus F-test evaluated all three groups. Gene Ontology over-representation analysis used the 74 retained and successfully mapped proteins as the background universe.

#### SomaScan

The SomaScan case study used the vendor-processed v4.1 ADAT file from PAD000036.^12^ The selected CADRE cord-plasma ADAT contained 7,596 aptamers and 130 study samples. Sixteen samples flagged by the vendor-provided RowCheck field were excluded, leaving 114 samples for analysis. endor hybridization normalization, median-intensity normalization, plate scaling, calibration, and vendor quality-control procedures represented in the ADAT metadata were not repeated. One sample without small-for-gestational-age status was excluded, leaving 7,596 aptamers and 129 samples for detection assessment. A measurement was classified as detected when its RFU was greater than or equal to the corresponding aptamer-specific mean buffer value. Aptamers detected in at least 80% of samples were retained first, followed by samples detected for at least 95% of retained aptamers. This retained 7,569 aptamers and all 129 samples. The reported RFU values were preserved without masking or imputation and were log2 transformed before principal-component analysis. Cohort was specified as the batch variable, and small-for-gestational-age status was included as a protected biological variable during batch correction.

### Synthetic Performance Benchmarks

Synthetic matrices were generated by resampling protein rows and sample columns from the empirical MS example with replacement. Noise with a standard deviation of 0.2 was added on the log2 scale while retaining the empirical missingness pattern. Samples were assigned to balanced control and treatment groups, and a log2 fold change of 1 was added to a randomly selected 10% of proteins in treatment samples. Batch labels were crossed with treatment condition. Protein-axis benchmarks evaluated 1,000, 2,000, 4,000, 8,000, and 16,000 proteins with 32 samples. Sample-axis benchmarks evaluated 8, 16, 32, 64, 128, 256, 512, and 1,000 samples with 8,000 proteins.

Five replicates of each size were run on a 2021 Apple M1 Pro computer with 16 GB memory, macOS 15.6.1, R 4.6.0, reference BLAS, and one computational thread. Each package stage was executed in a fresh R process. Elapsed time and peak process resident-set size were measured for import, quality control, preprocessing, clustering, differential analysis, abundance visualization, and pathway analysis. Plot objects were rendered during timing. Sample-correlation benchmarks used the 1,000 highest-variance proteins; this benchmark setting differs from the current Shiny interface starting value of 1,000. Shiny latency was measured through automated browser sessions that included file upload and rendering of representative workflow pages. Pathway runtime was interpreted in relation to the number of selected genes rather than as a direct function of matrix dimensions. Complete operation definitions and raw measurements are provided in the Supporting Information.

## RESULTS

### Advance over ProtPipe and Related Software

ProtPipe2 extends the original ProtPipe by supporting additional proteomics data types and making each analysis step available as a reusable R function. The original ProtPipe combines primary data-independent acquisition processing with command-line execution, containerized dependencies, and downstream reporting.^3^ ProtPipe2 instead begins with processed protein-abundance data and provides dedicated import functions for generic mass spectrometry matrices, Olink NPX exports, and SomaScan ADAT files. Each import function produces a SummarizedExperiment object, allowing all three data types to be analyzed using the same functions for quality control, preprocessing, statistical analysis, pathway enrichment, and visualization.

Table 1 compares ProtPipe2 with related downstream proteomics tools. Perseus retains analysis parameters within a desktop workflow; amica permits processed results to be downloaded and reloaded; ProTIGY can export results, an R workspace, and an optional R Markdown report; and MSstatsShiny can generate an R script corresponding to the selected MSstats analysis.^3–7^ Among the evaluated tools, ProtPipe2 is distinguished by combining direct Olink and SomaScan import with reusable R functions and an interactive web application. The comparison evaluates input support, interface type, and reproducibility features. It does not compare statistical accuracy or computational performance. ProtPipe2 uses established methods, including limma. Its main contribution is to bring these methods into one workflow for mass spectrometry, Olink, and SomaScan data, rather than to introduce a new statistical test.

**Table 1.** Input-format support and interface availability among downstream proteomics tools evaluated using the versions available in September 2026.

| **Tool** | **MS input** | **Olink input** | **SomaScan input** | **Web-based** | **No-code** |
| --- | --- | --- | --- | --- | --- |
| ProtPipe2 | Yes | Yes | Yes | Yes | Yes |
| ProtPipe | Yes | No | No | Yes | No |
| Perseus | Yes | No | No | No | Yes |
| amica | Yes | No | No | Yes | Yes |
| ProTIGY | Yes | No | No | Yes | Yes |
| MSstatsShiny | Yes | No | No | Yes | Yes |
| ProteoSign v2 | Yes | No | No | Yes | Yes |
| DEP | Yes | No | No | No | No |
| OlinkAnalyze | No | Yes | No | No | No |
| SomaDataIO | No | No | Yes | No | No |

### Cross-Platform Case Studies Mass Spectrometry

All analyses shown in Figure 1 were performed on the same imported dataset without intermediate reformatting. Coefficient-of-variation distributions summarized within-time-point variability (Figure 1A), while Spearman correlation, hierarchical clustering, and principal-component analysis provided complementary views of sample similarity and global structure (Figure 1B–D). Targeted heatmaps showed the abundance patterns of selected stem-cell and neuronal markers across the seven time points (Figure 1E, F). Together, these analyses demonstrate that ProtPipe2 can carry a generic mass spectrometry abundance matrix from sample-level quality control to protein-level visualization within a single workflow.

**Figure 1.**
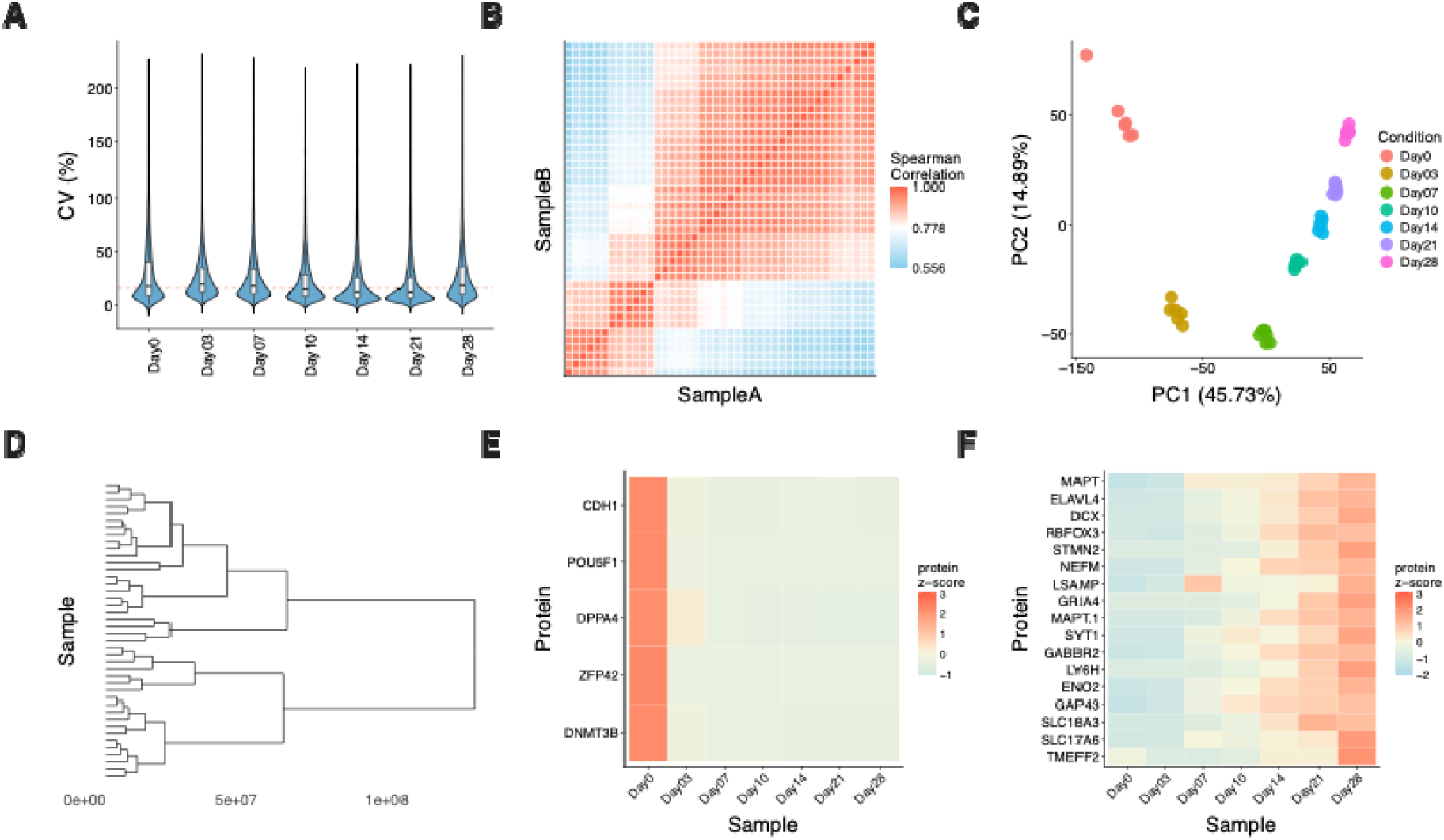
ProtPipe2 analysis of a generic mass spectrometry abundance matrix. **(A)** Coefficient-of-variation distributions across seven time points. **(B)** Spearman sample-correlation heatmap. **(C)** Principal-component analysis. **(D)** Hierarchical clustering. **(E, F)** Targeted heatmaps for stem-cell and neuronal marker sets. The input comprised 9,119 feature rows and 42 samples.

### Olink

mong the 92 assays and 126 samples remaining after sample-level Olink QC, 27 assays had at least one measurement below its plate-specific LOD or carrying an assay-level flag (Figure 2A). Applying the 80% assay-detection threshold retained 74 assays. Sample detection rates were then calculated across these assays, and 124 of 126 samples met the 95% threshold (Figure 2B). The retained dataset included 59 samples without diabetes, 22 with prediabetes, and 43 with type 2 diabetes.

**Figure 2.**
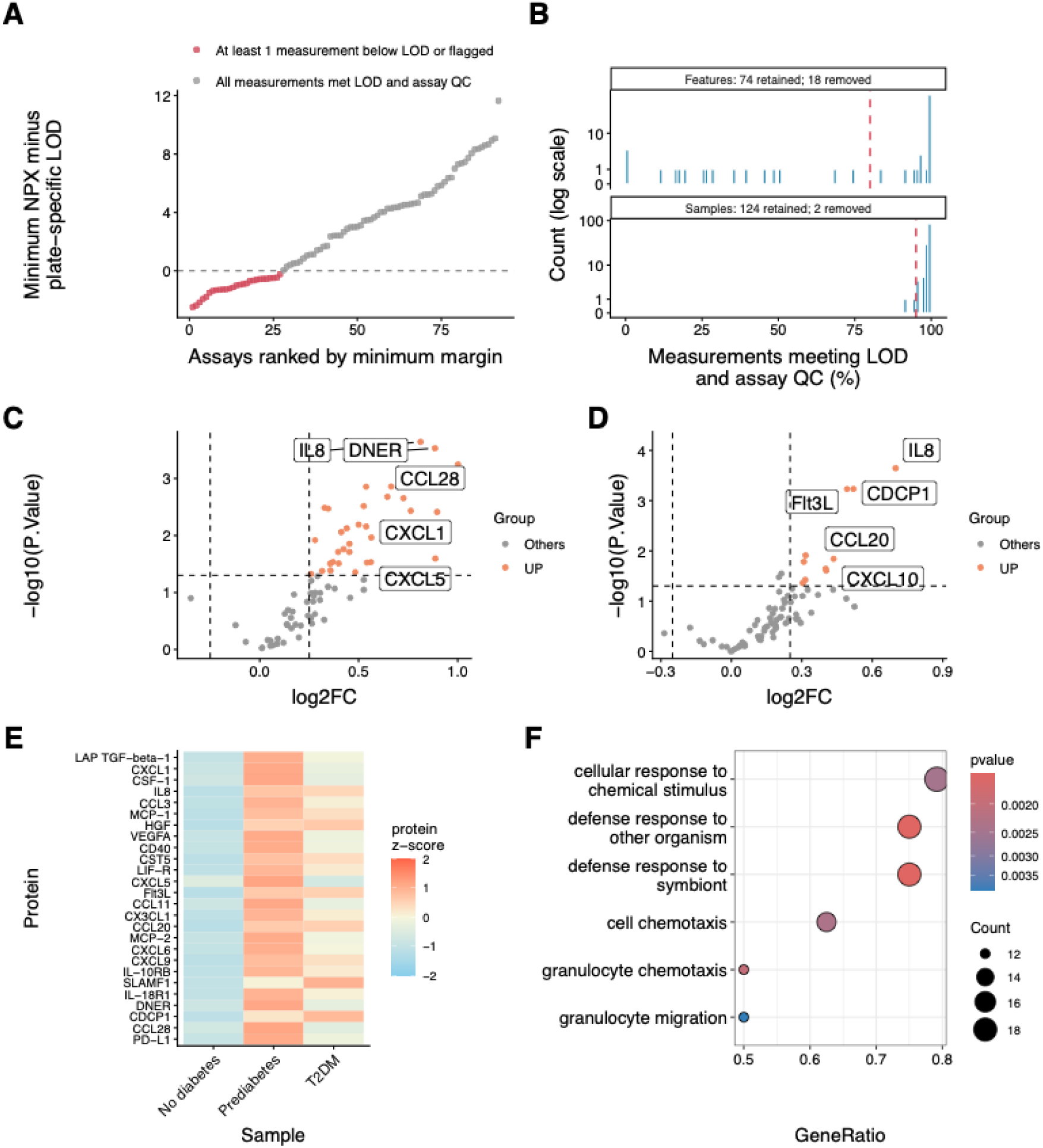
Olink PAD000040 detection filtering and downstream analysis. **(A)** Minimum NPX-minus-LOD margin for each assay, calculated using the plate-specific LOD of each measurement. Red points indicate assays with at least one below-LOD or assay-flagged measurement. **(B)** Assay- and sample-level detection-rate distributions. Dashed lines show the 80% assay threshold and 95% sample threshold. Seventy-four of 92 assays and 124 of 126 QC-passing samples were retained. Volcano plots compare **(C)** prediabetes and **(D)** type 2 diabetes with no diabetes; orange points meet nominal P ≤ 0.05 and |log2 fold change| ≥ 0.25. **(E)** Group-mean z-score heatmap for 26 assays meeting nominal three-group moderated-F P ≤ 0.05. **(F)** Six strongest nominal Gene Ontology Biological Process terms obtained using the 74 retained proteins as the background universe. No displayed term survived Benjamini–Hochberg correction.

Using the retained NPX values without imputation, 32 assays met the nominal visualization criteria of P ≤ 0.05 and |log2 fold change| ≥ 0.25 for prediabetes versus no diabetes, and 10 met these criteria for type 2 diabetes versus no diabetes (Figure 2C,D). The three-group moderated F-test identified 26 assays at nominal P ≤ 0.05, of which five remained significant after Benjamini–Hochberg adjustment. The group-mean heatmap shows the abundance patterns of the 26 nominally selected assays (Figure 2E). Gene Ontology analysis used the 74 retained proteins as the measured background and the 26 nominally selected proteins as the foreground (Figure 2F). No tested term remained significant after Benjamini–Hochberg adjustment; therefore, the enrichment panel demonstrates the analysis and background-selection workflow rather than providing evidence of pathway enrichment.

### SomaScan

The SomaScan case study demonstrates buffer-based detection filtering without replacing or imputing the reported RFU values (Figure 3). Comparison of each aptamer’s minimum sample signal with its buffer value identified aptamers with at least one measurement below the buffer reference (Figure 3A). At the selected thresholds, 7,569 of 7,596 aptamers met the 80% feature-detection criterion, and all 129 samples met the 95% sample-detection criterion after feature filtering (Figure 3B).

**Figure 3.**
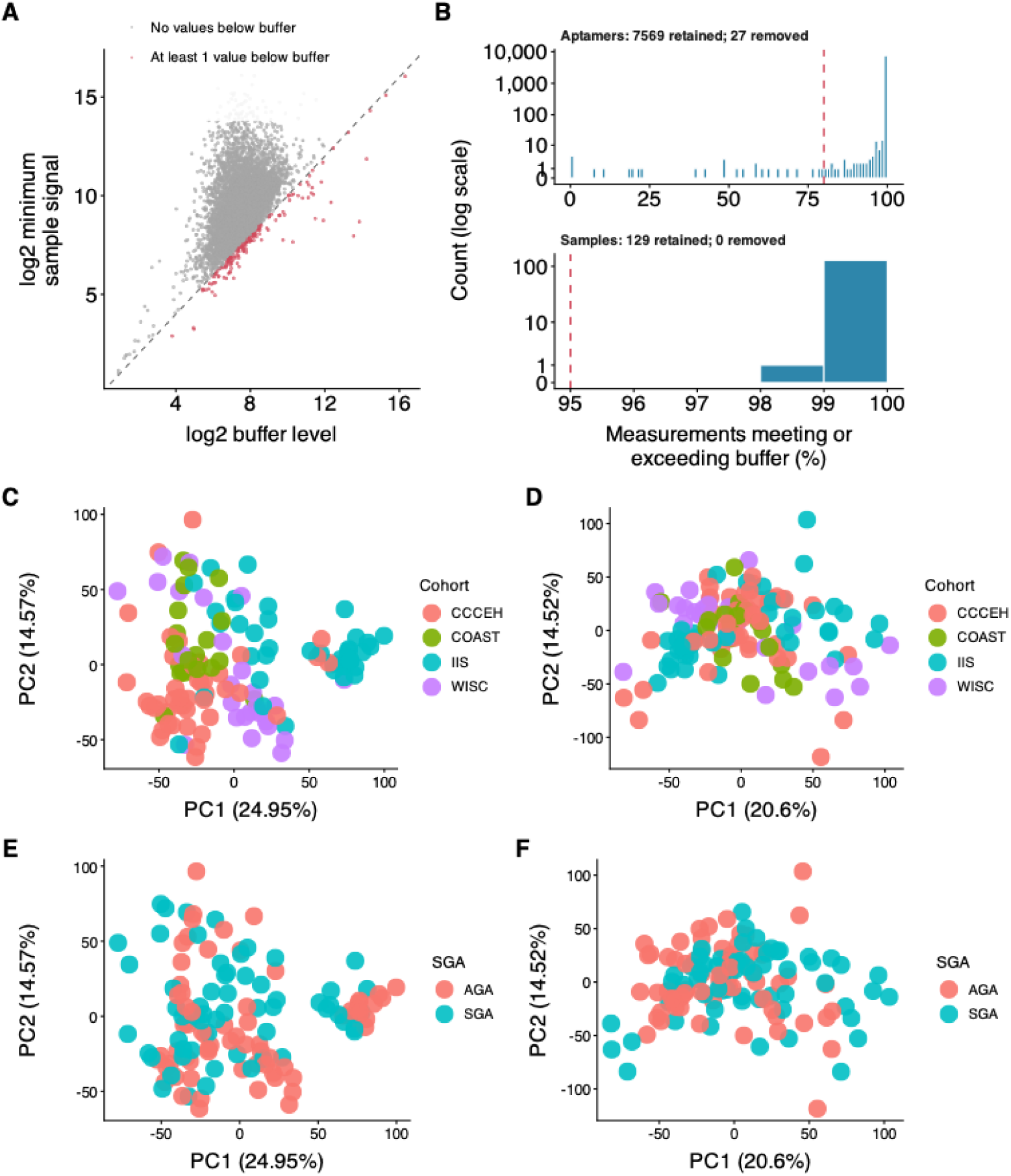
Buffer-based detection filtering and cohort correction of the SomaScan dataset. **(A)** Minimum sample signal versus the feature-specific buffer level. Highlighted features have at least one measurement below the buffer level and therefore fall below the dashed identity line. **(B)** Distributions of aptamer- and sample-level detection rates. Aptamer-level detection was defined as the percentage of samples in which the signal met or exceeded the corresponding buffer level. Sample-level detection was defined as the percentage of aptamers for which the signal met or exceeded the corresponding buffer level. Dashed vertical lines indicate filtering thresholds of 80% for aptamers and 95% for samples. All 129 samples were retained; 7,569 of 7,596 aptamers were retained and 27 were removed. Counts are shown on a logarithmic scale. **(C–D)** Principal component analysis colored by cohort before and after cohort correction, respectively. **(E–F)** The same principal component analyses colored by small-for-gestational- age status before and after cohort correction, respectively.

Principal-component analysis of the log2-transformed retained values showed cohort-associated structure before correction (Figure 3C). This separation was reduced after cohort correction (Figure 3D). Small-for-gestational-age status was protected during correction and is shown before and after correction in Figure 3E,F. These results demonstrate buffer-based detection assessment, ordered feature and sample filtering, and protected-variable batch correction. They do not establish that the displayed thresholds or correction model are appropriate for every SomaScan study.

### Computational Scaling

With the sample count fixed at 32, runtime increased modestly as the number of proteins increased for most package operations (Figure 4A). With the protein count fixed at 8,000, increasing the number of samples had the greatest effect on clustering and quality-control runtime (Figure 4B). For the largest matrix tested (8,000 proteins × 1,000 samples), clustering was the slowest operation, with a median runtime of 80.1 s. Quality control had the highest peak memory use at 2.84 GiB, compared with 0.608 GiB for the dependency-loaded R baseline (Figure 4C, D). Memory use for all tested operations remained below the 8 GiB reference line. In the interactive web application, the longest observed response time was 17.7 s for clustering the 8,000 × 1,000 matrix (Figure 4E, F). These measurements describe ProtPipe2 performance on the tested computer and should not be interpreted as a direct performance comparison with other software.

**Figure 4.**
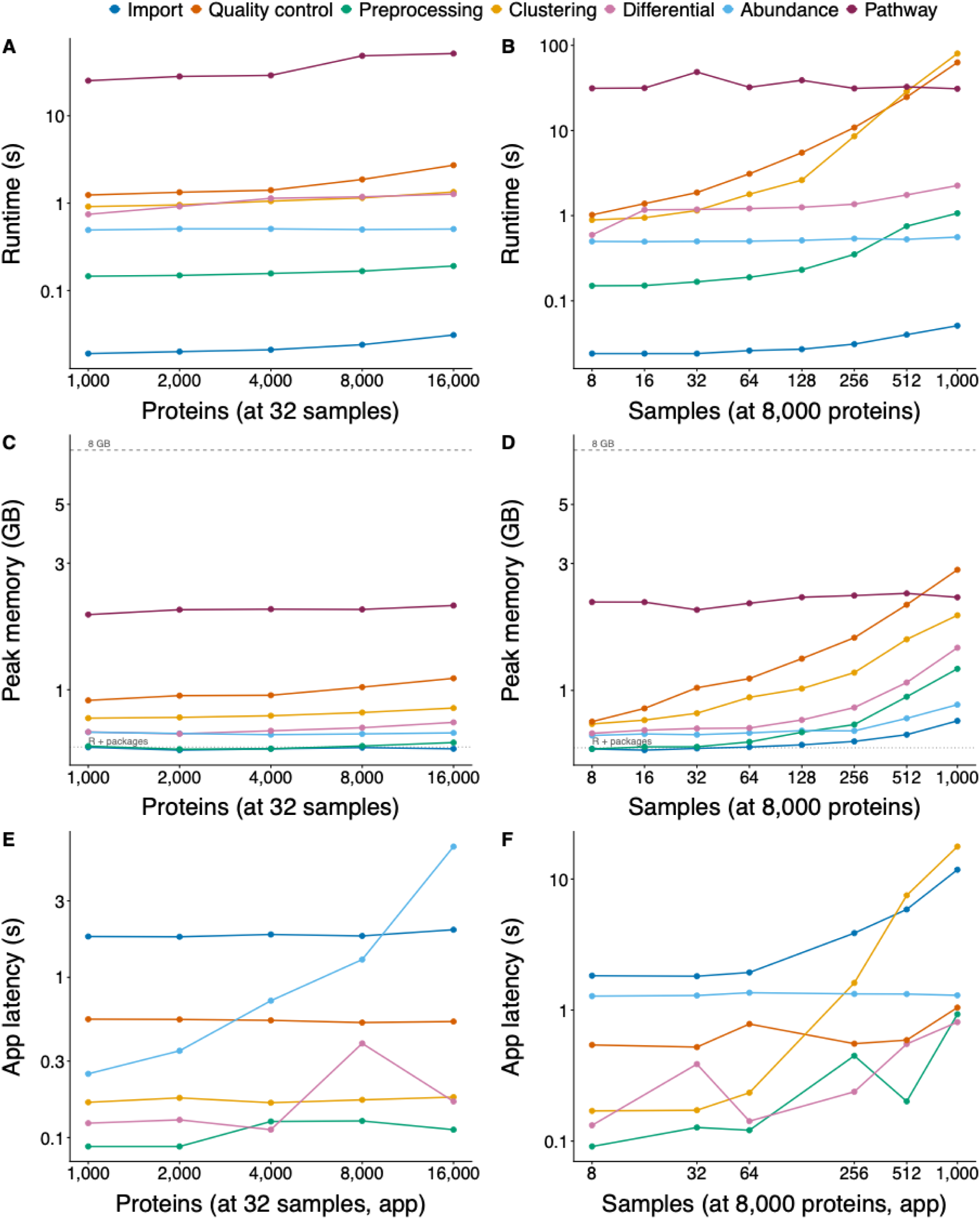
Synthetic scaling benchmarks with five replicates per size. Package median runtime versus (A) proteins and (B) samples; peak resident memory versus (C) proteins and (D) samples; Shiny operation latency versus (E) proteins and (F) samples. Numeric labels in A and B are log-log slopes. Pathway runtime depends on the selected-gene set, and synthetic enrichment results have no biological interpretation.

## DISCUSSION

ProtPipe2 provides a common downstream analysis workflow for mass spectrometry, Olink, and SomaScan data through an R package and an interactive web application. Unlike the original ProtPipe, which combines primary DIA processing with downstream reporting, ProtPipe2 begins with processed protein-abundance data and supports all three proteomics platform. Its main advantage is that the same analysis functions support both graphical and scripted workflows. Users can therefore perform an analysis in the web application and then inspect, repeat, or extend it in R.

The three case studies show how ProtPipe2 applies a common workflow while retaining information specific to each platform. The MS workflow demonstrates quality-control visualization and downstream exploration of a generic abundance matrix. The Olink workflow incorporates assay- and sample-level quality flags and measurements below the supplied limits of detection before downstream statistical analysis. The SomaScan workflow preserves ADAT annotations and masked values while supporting optional cohort correction and visualization. These examples demonstrate how the software operates and are not intended to reproduce the biological conclusions of the source studies.

Using the same functions in the R package and web application provides a direct path from interactive exploration to reproducible analysis. Processing choices are recorded with the data and can be exported with the results. Software tests cover data import, metadata handling, platform-specific quality control, statistical calculations, and plot generation. Synthetic benchmarks characterize changes in runtime and memory use as the numbers of proteins and samples increase. ProtPipe2 completed the tested workflows for matrices containing up to 8,000 proteins and 1,000 samples on the tested system, although performance depends on the selected analysis and computer configuration.

ProtPipe2 has several analytical limitations. It begins with processed abundance measurements and does not replace platform-specific primary processing, calibration, or vendor quality control. It also does not automatically determine the appropriate missingness model, normalization procedure, covariates, batch definition, statistical contrast, background set for pathway analysis, or reporting threshold. These decisions remain dependent on the experimental design and measurement platform. In particular, batch correction may remove biological variation when technical and biological factors are confounded, while imputation may influence downstream effect estimates. Associations with continuous variables should therefore be interpreted in the context of the study design and should not be considered evidence of causation.

The demonstrations and benchmarks also have limitations. The case studies were selected to demonstrate the three supported input types and do not provide biological ground truth for comparing statistical accuracy. The synthetic matrices were derived from one empirical MS dataset and therefore may not reproduce all characteristics of Olink or SomaScan data. Benchmarks were conducted on one computer with one R configuration and did not evaluate other operating systems, multiple simultaneous web-application users, server capacity, or network conditions. In addition, the privacy and performance of the web application depend on its deployment environment; restricted or regulated data should instead be analyzed using the R package or a locally deployed application under institutionally approved controls. Future development may extend the supported input formats, statistical models, and reporting functions while retaining a shared implementation across the R package and web application. Within these limits, ProtPipe2 provides an accessible workflow while keeping the analysis steps available for inspection and reproduction.

## CONCLUSION

ProtPipe2 provides a common workflow for the downstream analysis of mass spectrometry, Olink, and SomaScan proteomics data. By providing both an R package and an interactive web application, ProtPipe2 supports graphical and scripted analysis using the same underlying functions. The case studies demonstrate the three supported workflows, while the benchmarks characterize computational performance across datasets of different sizes. ProtPipe2 extends the original ProtPipe into an accessible and reproducible tool while leaving study-specific analytical decisions under user control.

## ASSOCIATED CONTENT

### Data and Software Availability

Source code is available at https://github.com/NIH-CARD/ProtPipe2 and documentation at https://nih-card.github.io/ProtPipe2.

## AUTHOR INFORMATION

### Corresponding Authors

Ziyi Li - Center for Alzheimer’s and Related Dementias, National Institute on Aging and National Institute of Neurological Disorders and Stroke, National Institutes of Health, Bethesda, Maryland 20892, United States; DataTecnica LLC, Washington, DC 20037, United States.

Yue Andy Qi - Center for Alzheimer’s and Related Dementias, National Institute on Aging and National Institute of Neurological Disorders and Stroke, National Institutes of Health, Bethesda, Maryland 20892, United States.

### Author Contributions

Conceptualization, J.E., C.W., Z.L., and Y.A.Q.; Software, J.E. and C.T.; Formal Analysis, J.E.; Validation, I.K., Y.H., B.J., and Z.L.; Data Curation, Y.H.; Visualization, J.E.; Supervision, M.R.C., M.A.N., Z.L., and Y.A.Q.; Writing – Original Draft, J.E. and Z.L.; Writing – Review and Editing, C.W., I.K., Y.H., C.T., B.J., M.R.C., M.A.N., Z.L., and Y.A.Q

## ACKNOWLEDGMENTS

This research was supported in part by the Intramural Research Program of the National Institutes of Health, National Institute on Aging, project ZIAAG000535. The contributions of the NIH author(s) are considered Works of the United States Government. The findings and conclusions presented in this paper are those of the author(s) and do not necessarily reflect the views of the NIH or the U.S. Department of Health and Human Services. All other authors declare no competing interests.

## Figures

**Supplementary Table 1.** ProtPipe2 implementation defaults and initial Shiny settings.

| Step | Parameter | Default value when selected | Use and interpretation |
| --- | --- | --- | --- |
| Minimum-intensity filtering | Minimum intensity | 0 | Available for MS data. Values below the selected threshold are treated as missing. |
| Filtering | Minimum protein detection | 80% | Retains proteins detected in at least the selected percentage of samples. |
|  | Minimum sample detection | 95% | Retains samples meeting the selected detection rate after protein filtering. |
|  | Filtering order | Proteins first, followed by samples | Sample detection rates are recalculated using the retained proteins. |
| Transformation | Transformation | $\log_2(x + 1)$ | Applied only when selected and appropriate for the input scale. Olink NPX is already log2-scaled. |
| Normalization | Normalization method | Median | Mean normalization may also be selected. |
| Imputation | Imputation method | Selected by the user | Available methods are listed below. No imputation is recommended by default for Olink or SomaScan. |
|  | Left-shifted distribution | Shift = 1.8; width = 0.3 | Suggested for MS data when missingness is expected to reflect low protein abundance. |
|  | K-nearest neighbors | k = 10 | Estimates missing values using similar protein profiles. |
|  | Random forest | 100 trees; 10 iterations | May require substantial runtime and memory for large datasets. |
|  | Protein-wise mean or median | No additional parameter | Uses the observed mean or median of the corresponding protein. |
|  | Scaled protein minimum | Multiplier = 1 | Uses the minimum observed value for each protein. |
|  | Fixed value | Value = 0 | Uses a user-defined constant. |
| Batch correction | Batch variable | Selected by the user | Selected from the available sample metadata. |
|  | Biological variables to protect | Selected by the user | Preserves variation associated with the selected biological variables. |
| Processing history | Recording | Automatic after an operation is selected | Records the operation and its parameter values. |
| Downloads | Processed data | TSV | Exports the processed abundance matrix. |
|  | Processing report | Markdown | Exports the processing history and parameter values. |

